# Quick, Don’t Move!: Wh-Movement and Wh-In-Situ Structures in Rapid Parallel Reading – EEG studies in English, Urdu, and Mandarin Chinese

**DOI:** 10.1101/2024.05.06.592830

**Authors:** Dustin A. Chacón, Donald Dunagan, Jill McLendon, Hareem Khokhar, Zahin Hoque

## Abstract

A fundamental question in the cognitive neuroscience of language is how grammatical representations are reflected in the organization and activity of the brain. This is challenging in part because superficial differences between languages, e.g., word order, exert different demands on the memory systems that process these structures. Here, we present an electroencephalography (EEG) study to investigate the brain’s responses to *wh*-constructions in English, Urdu, and Mandarin Chinese. In addition to the different word orders, writing systems, and morphological typologies of the 3 languages, these languages diverge with regard to how *wh*-constructions are formed: English requires filler-gap dependencies for *wh*-objects, whereas Urdu and Mandarin Chinese do not. We use a rapid parallel reading task, in which short sentences are displayed in parallel for 200ms to mitigate the different demands placed on memory systems. We show that neural responses distinguish *wh*-object constructions from their controls in midline anterior (Urdu, Mandarin Chinese) and right posterior sensors (English, Mandarin Chinese), from 200–400ms (English, Mandarin Chinese) and 500–800ms (English, Urdu). Although there is no detectable uniform, language-invariant response to *wh*-constructions across languages, there are a number of shared features in the evoked response between any pair of languages, i.e., *wh-in-situ* constructions generate an evoked response in midline anterior sensors. Moreover, behavioral evidence shows a robust cross-language cost of processing *wh*-object constructions, regardless of their surface form. This demonstrates that readers of diverse languages can process some grammatical information in a short 200ms fixation, and that the RPVP methodology may enable new ways of linking cognitive neuroscience of language to comparative syntax, i.e., the systematic description of similarities and differences between grammatical structures.

## 1. Introduction

A key question in the cognitive neuroscience of language is how grammatical structure is represented in the brain. This is a hard question. Experimental tasks in language comprehension require participants to process some word or sentence in speech, reading, or other modality. The temporal properties of the modality engage the attention, memory, and prediction networks in different ways, making it difficult to disentangle which brain responses correspond to grammatical representations vs. which correspond to the memory and attention mechanisms necessary for processing the stimulus. This problem is harder given the wide range of linguistic diversity. Across languages, comprehenders must manipulate different information at different times, depending on the grammatical properties of the language (see Chacón et al. 2024). Nonetheless, a relatively uniform network of left fronto-temporal and temporo-parietal areas appear to support language processing across different languages (Malik-Moraleda et al. 2022), and careful cross-language comparison of different languages has proven to be fruitful for understanding how the universal relates to the language-particular (e.g., Bornkessel-Schlesewsky & Schlesewsky 2016).

Our research question is: Is it possible to identify a single uniform neural response to a complex syntactic/semantic structure across languages, even if that structure may surface differently? Here, we present three experiments on the processing of simple main clause questions in English, Mandarin Chinese, and Urdu. All three experiments used the rapid parallel visual presentation paradigm (RPVP; Snell & Grainger 2017; Snell & Grainger 2019; Fallon & Pylkkänen 2024; Flower & Pylkkänen 2024; Krogh & Pylkkänen 2024; Dunagan et al. 2024; Dufau et al. 2024) with electroencephalography (EEG) recordings. In all three languages, participants read polar (‘yes/no’) questions, questions with *wh*-objects, questions with *wh*-subjects, and questions with both *wh*-subjects and *wh*-objects. If we permit participants to access the entire sentence structure in parallel, can we identify a uniform neural response corresponding to *wh*-constructions?

*Wh*-constructions are a particularly good test-case for investigating the mapping between variation in grammar, processing, and brain activity. Across languages, *wh*-constructions appear superficially different. For instance, English places (‘moves’) *wh*-phrases to the left edge of the clause in which they take semantic scope. In (1), the *wh*-object *who* is associated with a ‘gap’, indicated by _____, indexing the usual position that object NPs appear in English. By contrast, in Mandarin Chinese and Urdu, *wh*-phrases are *in-situ*, i.e., they do not ‘move’ and instead surface in the usual thematic position. Thus, the *wh*-object appears after the verb in Mandarin, a subject-verb-object language, (2), and before the verb in Urdu, (3), a subject-object-verb language. Despite this superficial difference, research in syntax and semantics has suggested that these structures share many representational features. Languages like Mandarin Chinese, for instance, are often analyzed as containing a relation between the *wh*-phrase and the left edge of the clause (‘Spec,CP’ Huang 1982; Beck 2006; Cheng 2009; Kotek 2019, among many others), which is motivated by the semantic scope of the *wh*-phrase. In (2–3), this ‘gap’ is marked with ______. Finally, setting aside the superficial details of the surface syntax, the sentences in (1)–(3) should be assigned a (virtually) identical semantic representation. For instance, if the semantics of *wh*-questions denote the set of possible answers to the question (4; Hamblin 1973, Karttunen 1977, Beck 2006, Kotek 2019), then the sentences in (1–3) elicit the same sets of answers. Thus, we refine our research question to: what brain activity is in common between the superficially different structures in (1)–(3) – if any?

1. [_CP_ **Who** did [_TP_ John see ____ ?]]
2. 约翰看了谁? [_CP_ [_TP_ Yuēhàn kàn-le **shéi ?**]] John saw who?
3. یونسنےکسکودیکھا؟ [_CP_ [_TP_ Yūnus-ne **kis-ko** dekhā ?]] John-Erg **who-Acc** saw.3Sg
4. ⟦Who did John see? ⟧ = { John saw *Mary*, John saw *Tom*, John saw *Chris*, …}

### 1.1 Representation and Processing of *Wh*-Dependencies

In psycholinguistic research, significant attention has been paid to how readers of languages like English process filler-gap dependencies, i.e., the relation between a moved *wh*-phrase and its gap. Overwhelmingly, findings demonstrate that filler-gap dependencies are processed ‘actively’ – readers attempt to relate a *wh*-phrase with a verb, even before the gap is confirmed. This pattern is observed in English (Crain & Fodor 1985; Stowe 1986; Traxler & Pickering 1996; Phillips 2006; Omaki et al. 2015), Japanese (Aoshima et al. 2004; Omaki et al. 2014), French (Bourdages 1992), Dutch (Frazier & Flores d’Arcais 1989), Bengali (Chacón et al. 2016), Urdu/Hindi (Dash & Kaiser 2019), Chamorro (Wagers et al. 2015) and Tagalog (Pizarro-Guevara & Wagers 2020). After the gap is encountered, readers may then need to retrieve the *wh*-phrase from memory using a cue-based retrieval mechanism to fully interpret it (McElree et al. 2003; Wagers & Phillips 2014).

Neural evidence also demonstrates comprehenders’ eagerness to interpret *wh*-constructions early, with early event-related potential (ERP) responses to unexpected or semantically unusual *wh*-verb relations (Garnsey et al. 1989; Dallas et al. 2013; Jessen & Felser 2019; Chacón & Pylkkänen 2024). Before the gap is encountered, filler-gap dependencies also elicit a sustained anterior negativity (SAN) from the onset of the *wh*-phrase, persisting until the gap is encountered or after (King & Kutas 1995; Kaan et al. 2000; Fiebach et al. 2002; Phillips et al. 2005). The SAN has been argued to reflect the neural correlate of maintaining the *wh*-phrase in working memory (Wanner & Maratsos 1978), although later work has questioned this interpretation (Yano & Koizumi 2019, 2021; Lo & Brennan 2021; Cruz Heredia et al. 2022). A distinct neural signature is also observed after the gap is encountered in serial reading, usually engendering a ‘P600’ response, a positive-going ERP located over centro-parietal and posterior sensors, from around 500–900ms post-verb onset (Kaan et al 2000; Phillips et al. 2005). This P600 response may reflect retrieval and integration of the *wh*-phrase. Crucially, these behavioral responses and ERP responses are interpreted through the lens of the memory computations needed to relate the *wh*-phrase and the gap while processing a sentence word-by-word in serial reading tasks.

Left inferior frontal gyrus (LIFG; ‘Broca’s Area’) appears to be systematically implicated in processing *wh*-constructions (Just et al. 1996; Stromsworld et al. 1996). This activity may reflect the representation of movement relations (Ben-Shachar et al. 2003, 2004), irrespective of modality. Alternatively, LIFG activity may correlate with more general cognitive operations deployed in language processing, such as memory retrieval (Just & Carpenter 1992; Smith & Jonides 1999; Rogalsky et al. 2008; Leiken & Pylkkänen 2014) or cognitive control (Miller & Cohen 2001; Novick et al. 2005; Braver 2012). Some evidence suggests a mixed view, in which LIFG activity is modulated by both movement relations and working memory demand independently (Santi & Grodzinsky 2007; Leiken et al. 2015). This again underscores the challenge in dissociating neural responses corresponding to the representation of a grammatical structure vs. the cognitive systems needed to process and interpret it.

Less attention has been paid to the processing of *wh-in-situ* phenomena, as observed in Mandarin Chinese or Urdu. However, serial reading results demonstrate that Mandarin Chinese readers must retrieve the boundary of a scope-taking clause when encountering a *wh*-element (the ‘Spec,CP’ position; Xiang et al. 2014, 2015; see also Kraft et al. 2018 and Yang et al. 2023). These results suggest that even readers of *wh-in-situ* languages like Mandarin Chinese must deploy a memory retrieval strategy cued at the terminal element of a *wh*-construction when reading a sentence word-by-word, much like English readers must at a gap site. To our knowledge, there have been no studies examining the processing of simple *wh*-*in-situ* constructions in Urdu/Hindi (however see Dash & Kaiser 2019 on long-distance scope-marking structures in Hindi).

### 1.2 Rapid Parallel Visual Presentation

To our knowledge, most theories of processing *wh*-constructions have focused on relating the two elements of the dependency (the *wh*-phrase and the gap or clause edge) over time, while reading or hearing a sentence word-by-word. However, it is unclear what these models predict about the processing of *wh*-constructions when the elements of the dependency are not separated temporally. We address this using RPVP – a whole-sentence, single-fixation reading paradigm. Participants fixate on a single sentence presented in the foveal center for a short duration, usually 200ms, followed by a memory probe. Previous results demonstrate that comprehenders are capable of extracting some sentence-level syntactic and semantic features of the sentence in a single fixation in Japanese (Asano et al. 2011), French (Snell & Grainger 2017, 2019; Wen et al. 2019; Dufau et al. 2024), English (Dunagan et al. 2024; Fallon & Pylkkänen 2024, Flower & Pylkkänen 2024) and Danish (Krogh & Pylkkänen 2024). This presentation style mitigates the burden placed on the memory system by more standard word-by-word presentation styles used in cognitive neuroscience of language experiments. Superficial details, such as basic word order, argument-verb agreement, and *wh*-movement, may impact working memory resources less than in serial reading studies. This is because all members of relevant grammatical dependencies, e.g., the *wh*-word and the verb, are presented to the visual system at the same time, mitigating the need to maintain and retrieve information throughout the duration of the sentence. We propose that any commonality observed in evoked neural responses to grammatical structures in an RPVP task could index the brain’s uniform representation of that structure, abstracting away from the details of navigating word-by-word processing.

Using a similar logic, Krogh & Pylkkänen (2024) used an RPVP task to investigate the magnetoencephalography (MEG) response to argument structure alternatives and yes/no questions in Danish, both modeled using verb movement. They identified a neural response localized in right fronto-medial regions ∼260–370ms corresponding to argument structure alternations, followed by a different pattern in right occipital areas ∼500–550ms and parietal areas ∼640–720ms. This demonstrates that processing mechanisms deployed in RPVP are sensitive to the syntactic and semantic properties of verb movement in Danish. However, these alternations are ‘local,’ i.e., correspond to the position of the verb in simple main-clause sentences, which are not commonly studied in psycholinguistics of filler-gap relations, and may not undergo the same kinds of processing dynamics as *wh*-constructions in English. Additionally, Krogh & Pylkkänen (2024) only compared different sentence structures within the same language, because their aim was to establish the brain’s sensitivity to these grammatical alternations in Danish. By contrast, our study seeks to identify the commonalities and differences in the neural response to superficially different *wh*-constructions in English, Urdu, and Mandarin Chinese.

### 1.3 Comparing the Syntax of English, Urdu, and Mandarin *Wh*-Constructions

We chose English, Mandarin Chinese, and Urdu because of their many superficial differences. Any commonality in the EEG response to *wh*-constructions across these languages could identify a putative neural response corresponding to the shared syntactic or semantic features of these sentences. English is a predominantly subject-verb-object language with some flexible word order, with fusional morphology, subject-verb agreement, and is predominantly written left-to-right in the Roman script. Mandarin Chinese is a subject-verb-object language with strict word order, isolating morphology, no verbal agreement, and is predominantly written left-to-right in Chinese characters (Hanzi), but may be written vertically and right-to-left in more traditional texts. Urdu is a subject-object-verb language with flexible word order, complex fusional morphology, split-ergative agreement in which the verb agrees with the subject or object depending on case morphology, and is written right-to-left using the Nastaʿlīq variant of the Perso-Arabic script. Thus, the visual reflex of a short transitive sentence in these 3 languages is quite distinct with different information encoded in different positions of the sentence, which may engage with early reading processes differently.

The *wh*-constructions of these languages exhibit key syntactic differences. As discussed earlier, English overtly employs a filler-gap structure with *wh*-phrases, shown in (1) and repeated in (5). This grammatical rule applies even across multiple clauses, as seen with a long-distance *wh*-object question in (6) and a long-distance *wh*-subject question in (7). Moreover, English requires that a main clause containing a moved *wh*-phrase include an auxiliary (*do*-insertion), realized in (5–8) as the past tense *did*. This also includes polar (‘yes/no’) questions, (8).

Importantly, however, main clause *wh*-subjects do not exhibit this pattern. No *do*-insertion occurs in (9), and the subject *wh*-question *who* remains in its canonical position (see Pesetsky & Torrego 2001). Moreover, *wh*-subjects ‘block’ movement of *wh*-objects; sentences with *wh*-subjects and *wh*-objects must leave both in their canonical position, (10).

(5) [_CP_ **Who** did [_TP_ John see]] ?
(6) [_CP_ **Who** did [_TP_ Mary say [_CP_ [_TP_ John saw]] ?
(7) [_CP_ **Who** did [_TP_ Mary say [_CP_ [_TP_ saw John]] ?
(8) [_CP_ **Did** [_TP_ John see Mary?]]
(9) [_CP_ [_TP_ **Who** saw Mary]?
(10) [_CP_ [_TP_ **Who** saw **who**]?

Mandarin Chinese, by contrast, systematically leaves *wh*-questions in their canonical position. As seen in (2), repeated in (11), *wh*-objects surface in the post-verbal object position. This is true even in long-distance questions, (12). Mandarin Chinese uses a dedicated question marker 吗 *ma* to indicate a polar question, represented as Q in (13). Finally, as represented in (14), post-verbal *wh*-questions are ambiguous between a *wh*-interpretation and an indefinite variable interpretation (‘anyone’ / ‘someone’, Huang 1982; Dong 2009), demonstrating an ambiguity in the mapping between surface form and semantic structure. This interpretation is restricted to certain syntactic contexts.

(11) 约翰看了谁? [_CP_ [_TP_ Yuēhàn kàn-le **shéi**]] ? John saw who ‘Who did John see?’
(12) 玛丽说约翰看了谁? [_CP_ [_TP_ Mǎlì shuō [_CP_ [_TP_ Yuēhàn kàn-le **shéi**]] ? Mary said John saw who ‘Who did Mary say that John saw?’
(13) 约翰看了玛丽吗? [_CP_ [_TP_ Yuēhàn kàn-le Mǎlì **ma**]] ? John saw Mary Q
(14) 谁看了谁? [_CP_ [_TP_ **shéi** kàn-le **shéi**]] ? who saw who ‘Who saw who?’ ‘Who saw anyone?’

Urdu, like Mandarin Chinese, systematically leaves *wh*-questions in their canonical positions in main clauses, as shown in (3) and repeated in (15). For questions about embedded clauses, however, Urdu uses two distinct strategies. The *wh*-phrase may move the *wh*-phrase to the main clause, before the main verb and after the subject, shown in (16). More commonly, Urdu uses an ‘expletive *wh*’ in sentences with long-distance questions, exemplified in (17). Here, the word ی (*kyā* ‘what’ appears before the main verb to indicate the scope of the *wh-*question, and its correlated *wh*-phrase سکو *kis-ko* ‘who’ occurs in its thematic position. With this strategy, both elements of the *wh*-construction are overtly realized (Dayal 1996; Lahiri 2002). In addition to this scope-marking use, ی (*kyā* ‘what’ may also mark polar questions. The position of ی (*kyā* as a polar question marker is variable, but it typically occurs at the beginning of the clause, represented as Q in (18).

(15) یونسنےکسکودیکھا؟ [_CP_ [_TP_ Yūnus-ne **kis-ko** dekhā?]] John who saw? ‘Who did John see?’
(16) میرینےکسکوکہاکہیونسنےدیکھا؟ [_CP_ [_TP_ Mery-ne **kis-ko** kahā [_CP_ ki [_TP_ Yūnus-ne _____ dekhā]] ? Mary who said that John saw ‘Who did Mary say that John saw?’
(17) میرینےکیاکہاکہیونسنےکسکودیکھا؟ [_CP_ [_TP_ Mery-ne **kyā** kahā [_CP_ ki [_TP_ Yūnus-ne **kis-ko** dekhā]] ? Mary what said that John who saw ‘Who did Mary say that John saw?’
(18) کیایونسنےمیریکودیکھا؟ [_CP_ **kyā** [_TP_ Yūnus-ne Mery-ko dekhā]] ? Q John Mary saw ‘Did Mary see John?’

The details of the syntactic analysis of *wh*-constructions in English, Urdu and Mandarin Chinese remain controversial. The systematic positioning of *wh*-questions in their canonical position in Mandarin Chinese has motivated theories of ‘covert movement,’ analogizing the interpretation of moved and *in-situ wh*-phrases between English and Mandarin Chinese (Huang 1982; Cheng 2009). Other analyses instead argue for a more flexible mapping between *wh*-syntax and semantics (e.g., Tsai 1999; Erlewine 2024). Additionally, the different syntactic behavior of *wh*-phrases in multi-clausal structures in Urdu and Mandarin Chinese demonstrates that the same analysis cannot be applied to both languages without modification. Like Chinese, some authors have argued for covert movement in Urdu and Hindi (Mahajan 1990), an indirect association of the semantics of the *wh*-phrase and the scope marker ی (*kyā* ‘what’ (Dayal 1996; Lahiri 2002), or for ‘partial’ movement to a pre-verbal position (Spec,*v*P) as in (16) (Malhotra & Chandra 2007; Manetta 2010).

## 2. Methods

Our primary question is: Do English, Urdu, and Mandarin Chinese readers show distinct neural responses to *wh*-constructions vs. their controls in an RPVP task? If so, are there some common properties to this neural response which might reflect the shared syntactic or semantic structure of these sentences?

As a first blush hypothesis, we might expect a P600 response in all 3 languages, given that this is typically observed at the completion of a filler-gap response (Kaan et al 2000; Phillips et al. 2005), if the P600 reflects integration of a *wh*-phrase into the interpretation of a sentence. As suggested in Section 1.1, both filler-gap and *wh-in-situ* structures require memory retrieval operations in serial paradigms, and we therefore might expect similar responses across languages and reading paradigms as our null hypothesis.

However, if these P600 responses only correspond to integration of a filler phrase stored in memory during word-by-word processing, then we might not expect a typical P600. That is, if comprehenders can view both the elements of a *wh*-dependency simultaneously, they may not need to execute the same set of memory subroutines needed to maintain and retrieve a *wh*-phrase over time for overt filler-gap dependencies (English), or for accessing the left-edge scope position in memory (Urdu, Mandarin Chinese). If so, then neural responses may emerge much more quickly, i.e., on par with the 200–500ms findings observed by Krogh & Pylkkänen (2024).

### 2.1 Participants

Participants were recruited from the Athens, Georgia community. Participants must have been a self-identified native speaker of English, Mandarin Chinese, or Urdu. In total, there were 35 English participants, 33 Mandarin Chinese participants, and 27 Urdu participants. One participant from each language group was removed due to poor performance on the task (< 75% accuracy). Each participant’s language history was documented using the English, Urdu, and Mandarin Chinese versions of the Leap-Q questionnaire (Marian et al. 2007). Participants were also interviewed about their reading experience in the target language, to ensure that they had substantial facility in reading Roman, Chinese, or Nastaʿlīq script. Handedness was also tracked using the Edinburgh Handedness Survey (Oldfield 1971). *Post*-*hoc* analyses revealed no meaningful relation between handedness, language history, and task performance, and we did not detect any systematic differences between participants’ recorded EEG signals that could be explained by any of the demographic variables. Participants all gave written consent before participating, and were provided with consent forms in either the target language (English, Urdu, or Mandarin Chinese) or another language by request. Initial recruitment communications occurred in the target languages by native speaker experimenters.

### 2.2. Materials

For each language, we prepared 64 sets of stimuli. Each set consisted of 4 main-clause questions, distributed in a 2 × 2 design, manipulating ±Wh-Subject and ±Wh-Object.

The –Wh-Subject, –Wh-Object conditions were all polar questions. This was marked with the auxiliary *did* in English, and the Q particles ی (*kyā* ‘what’ in Urdu and 吗 *ma* in Mandarin Chinese.

The +Wh-Subject, –Wh-Object sentences included an indefinite *wh*-pronoun in the subject position: *who* in English, کسنے v *kis-ne* ‘who’ in Urdu, and 谁 *shéi* ‘who’ in Mandarin Chinese.

The –Wh-Subject, +Wh-Object sentences included a definite noun phrase with a *wh*-determiner in the object position: *which* <noun> in English, کونسا/کونسی *kaun sā/kaun sī* <noun> in Urdu, and 哪 *nǎ-*<classifier> <noun> in Mandarin Chinese. In the English sentences, the *wh*-object occurred at the left edge of the sentence, forming a filler-gap dependency, and was followed by *did*. Urdu requires agreement of the *wh*-determiner کونسا/کونسی *kaun sā/ kaun sī* in gender and number with the object noun, thus we used the masculine (ا – ā) and feminine (ی–ī) endings when appropriate. Mandarin Chinese requires a classifier that matches in semantic features with the noun it modifies for demonstratives like 哪 *nǎ* ‘which’ and 这 *zhè* ‘this,’ thus we chose the correct classifier for each noun.

For the +Wh-Subject, +Wh-Object conditions, all languages used the indefinite *wh*-pronoun in the subject position and the definite noun phrase with *wh*-determiner in the object position. In English, there was no filler-gap dependency or *do*-support.

To ensure similarity of word length and semantics across conditions and languages between +Wh and –Wh conditions, the non-*wh* were always indefinite pronouns (*she/he, us-ne*, 他/她 *tā*), and the non-*wh* objects were always definite noun phrases with proximate demonstratives (*this* <noun>, 这 ی *zhè*-<classifier> <noun>,,..,_ *yeh* <noun>).

All sentences were presented in the simple past in English and the present perfective in Urdu. In English. All verbs ended in *-ed* or a common irregular past tense form. In Urdu, all subjects were marked with the ergative case نے *-ne*, as required in the present perfective. Urdu uses a split-ergative agreement system, which requires that the verb agrees with the object in person, number, and gender in this tense/aspect combination. Thus, in all sentences, the verb agreed with the gender of the object noun. In Mandarin Chinese, we used the perfective particle了 *le* in all sentences, except for the polar question –Wh-Subject, –Wh-Object condition, which ended in the Q particle 吗 *ma*. This was meant to counter-balance the length of the stimuli, which seemed to be important to maintain equal comprehensibility across items in pilot studies with Mandarin Chinese-reading participants.

All sentences used singular or unmarked nouns only. Gender was counterbalanced across trials in all languages. In English, half of the stimuli used the masculine subject pronoun *he* in the –Wh-Subject conditions, and the other half used the feminine subject pronoun *she*. In Urdu, pronouns do not mark gender. However, all nouns are marked masculine or feminine. Thus, in half of the trials, we used a masculine object NP, and the other half used a feminine object NP, and we selected the appropriate form of کونسا/کونسی *kaun sā/ kaun sī* in the +Wh-Object conditions. In Mandarin Chinese, the subject pronoun *tā* is morphologically unmarked for gender, but the choice of Chinese character differs for a masculine referent vs. feminine referent. Half of the trials therefore used the masculine character 他 *tā* and half used the feminine character 她 *tā*.

English stimuli were presented in the sans serif Monaco font. Urdu and Mandarin Chinese stimuli were pre-rendered as image files prior to the experiment. Urdu stimuli were prerendered with the Noto Nastaliq font, and Mandarin Chinese stimuli were prerendered in simplified Chinese characters in the ST Heiti Light font.

An example set of the stimuli are presented with complete glosses in Table 1, and repeated in Figure 1A.

**Table 1.** Example set of stimuli used in experiment. English translations of Urdu and Mandarin Chinese stimuli are in the left column, and native orthography and gloss are in the right column. Urdu is written right-to-left, but glossed left-to-right. Abbreviations: Q = question marker; CL = classifier; ASP = aspect marker; ERG = ergative case; Fem = feminine gender-marking.

|  |  |  |
| --- | --- | --- |
| **English** | –Wh-Subject, –Wh-Object | *Did she read this book?* |
|  | +Wh-Subject, –Wh-Object | *Who read this book?* |
|  | –Wh-Subject, +Wh-Object | *Which book did she read?* |
|  | +Wh-Subject, +Wh-Object | *Who read which book?* |
| **Mandarin** | –Wh-Subject, –Wh-Object | 她看这本书吗? |
| **Chinese** | 'Did she read this book?' | she read this-CL<sub>book</sub> book Q? |
|  | +Wh-Subject, –Wh-Object | 谁看这本书了? |
|  | 'Who read this book?' | who read this-CL<sub>book</sub> book ASP? |
|  | –Wh-Subject, +Wh-Object | 她看哪本书了? |
|  | 'Which book did she read?' | she read which-CL<sub>book</sub> book ASP? |
|  | +Wh-Subject, +Wh-Object | 谁看哪本书了? |

**Table 1.** Example set of stimuli used in experiment.
|  | 'Who read which book?' | who read which-CL<sub>book</sub> book ASP? |
| --- | --- | --- |
| **Urdu** | *–Wh-Subject, –Wh-Object* | کیا اس نے یہ کتاب پڑھی؟ |
|  | 'Did she read this book?' | Q she-ERG this book<sub>Fem</sub> read<sub>Fem</sub> ? |
|  | *+Wh-Subject, –Wh-Object* | کس نے یہ کتاب پڑھی؟ |
|  | 'Who read this book?' | who-ERG this book<sub>Fem</sub> read<sub>Fem</sub> ? |
|  | *–Wh-Subject, +Wh-Object* | اس نے کون سی کتاب پڑھی؟ |
|  | 'Which book did she read?' | she-ERG which<sub>Fem</sub> book<sub>Fem</sub> read<sub>Fem</sub> ? |
|  | *+Wh-Subject, +Wh-Object* | کس نے کون سی کتاب پڑھی؟ |
|  | 'Who read which book?' | who-ERG which<sub>Fem</sub> book<sub>Fem</sub> read<sub>Fem</sub> ? |

**Figure 1.**
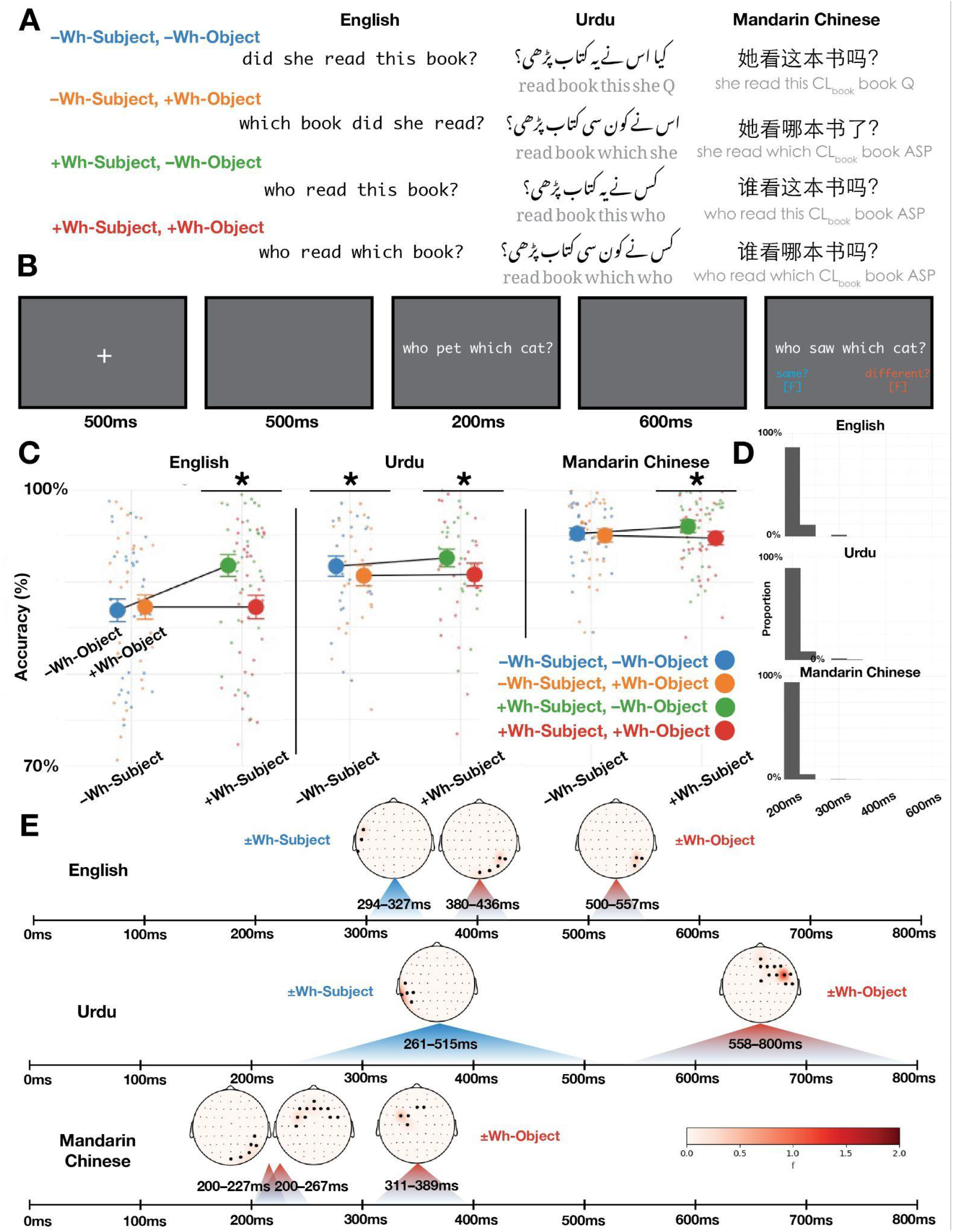
(A) Depiction of sample stimulus in presentation font in English, Urdu, and Mandarin Chinese. English word-by-word translations (glosses) are provided in gray below Urdu and Mandarin sentences. (B) Trial structure for RPVP experiment. (C) Accuracy on the memory probe by condition and language. Stars (*) indicate significant pairwise comparisons in the model described in Section 3.2. Error bars indicate one standard error from the mean above and below the mean. (D) Distribution of presentation times across all trials by language. (E) Summary of EEG results by language. Topoplots show the spatial distribution of significant clusters, and time spans printed below each cluster indicate temporal extent of the cluster. Blue time spans correspond to clusters in which +Wh-Subject trials had a greater evoked response (positive or negative) compared to –Wh-Subject trials; red time spans correspond to clusters in which +Wh-Object trials had a greater evoked response (positive or negative) compared to –Wh-Object trials.

### 2.3 Procedure

The RPVP experiment was conducted using PsychoPy. The procedure is adapted from the original RPVP paradigm introduced by Snell & Grainger (2017, 2019), and later elaborated by Fallon & Pylkkänen (2024), Krogh & Pylkkänen (2024), Flower & Pylkkänen (2024), and Dunagan et al. (2024). Stimuli were presented within-subjects, such that each participant saw each sentence. Order presentation was pseudo-randomized into 4 separate lists, such that one item set occurred in one condition per list, and each list contained an equal number of trials per condition. Between each list, participants took a break, in which participants read jokes or saw photos of baby animals.

Each trial consisted of a fixation cross, centered in the screen and displayed for 500ms, followed by a blank screen for 300ms. The critical sentence then appeared for a fixed duration, followed by another gray screen. Finally, an untimed probe sentence occurred. Participants were instructed to respond to whether the probe sentence was the same as the prior target sentence or different. Participants entered their response by pressing the [F] key to indicate that the sentences matched, and the [J] key to indicate that they mismatched. On-screen reminders were provided on the lower half of the screen during the probe sentence. Half of the trials changed the object noun or the verb to create ‘mismatch’ trials. An error message that lasted for 1000ms was provided as on-screen feedback for incorrect trials. The trial structure is exemplified in Figure 1B.

Because we are comparing performance across participants who read languages with substantially different writing systems, we followed the procedure in Dunagan et al. (2024) in dynamically adjusting the presentation time of the critical sentence. The critical sentence was initially displayed for 200ms with the subsequent blank screen displayed for 600ms, summing 800ms in total. If participants entered an incorrect response, then the time of the critical sentence increased by 50ms, and the blank screen display time decreased by 50ms. After a correct trial, the display time of the critical sentence decreased by 50ms, and the blank screen increased by 50ms. This dynamic adjustment of presentation time was constrained such that the critical sentence could not be faster than 200ms or slower than 600ms, and that the sum of the critical sentence and subsequent blank screen sum to 800ms. Faster display times are preferable for an RPVP task because it ensures that readers do not saccade, which would engender endogenous bioelectrical noise in the EEG recordings, and enable serial reading strategies.

Participants were kept at a uniform distance from the screen, approximately 70cm from nasion to center of the screen. The visual angle subtended of the stimuli was approximately 15 degrees. All stimuli were printed in a white font against a dark gray background in a dimly-lit room. All on-screen instructions and communications were conducted in the target language.

EEG recordings were done with a BrainVision® actiChamp+ amplifier. Participants were fitted with a 64 channel EEG cap, with Ag/Cl electrodes. Sensor impedances were kept below 20 kΩ using electro-conductive gel.

In addition to this procedure, each participant conducted an unrelated experiment, and participated in an unrelated localizer task. The short localizer task lasted approximately 5 minutes and was conducted first. The order of the two experimental tasks were counterbalanced. This experiment took approximately 20 minutes, and the whole procedure – including EEG preparation and cleanup – took approximately 90–120 minutes. Participants were compensated 10 USD per half hour of participation.

## 3. Results

### 3.1 Behavioral Results

We used the lme4 package in R to fit a logistic mixed effects regression model to the accuracy data. We fit the correct responses as the dependent variable, with factors ±Wh–Subject, ±Wh-Object, Language, and their interaction terms as predictors, and participant and item as random effects. In this model, –Wh-Subject and –Wh-Object were coded as 0, and +Wh-Subject and +Wh-Object were coded as 1. Language was contrast coded as two separate coefficients – English = (0, 1), Mandarin Chinese = (1, 0), and Urdu = (–1, –1). Afterwards, we conducted pairwise comparisons using the emmeans package, to compare the accuracy between each language group, and to compare each level of ±Wh-Object nested within each factor of ±Wh-Subject in each language. The results of the model are shown in Table 2, and the average accuracy by condition and language is shown in Figure 1C.

**Table 2.**
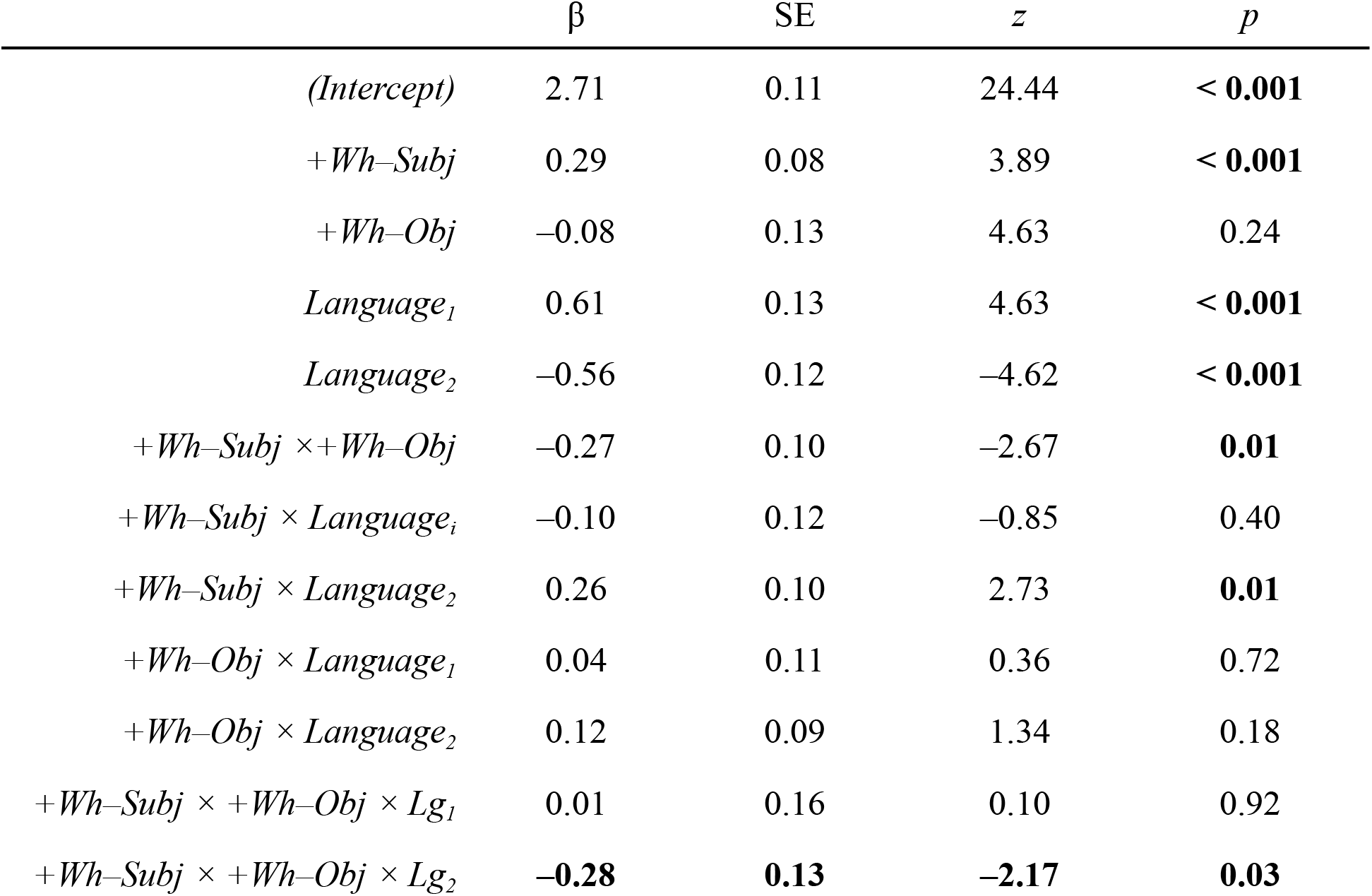
Results of logistic mixed effects model fit to the accuracy data. The structure of the model was Correct Responses ∼ ±Wh–Subject × ±Wh-Object × Language + (1|Participant) + (1|Item). ±Wh-Subject and ±Wh-Object were coded with –Wh conditions as 0 and +Wh conditions as 1; Language was contrast coded as two coefficients. Bolded *p*-values correspond to significant effects (*p* < 0.05).

| | $\beta$ | SE | $z$ | $p$ |
| --- | --- | --- | --- | --- |
| *(Intercept)* | 2.71 | 0.11 | 24.44 | **< 0.001** |
| *+Wh-Subj* | 0.29 | 0.08 | 3.89 | **< 0.001** |
| *+Wh-Obj* | -0.08 | 0.13 | 4.63 | 0.24 |
| *Language<sub>1</sub>* | 0.61 | 0.13 | 4.63 | **< 0.001** |
| *Language<sub>2</sub>* | -0.56 | 0.12 | -4.62 | **< 0.001** |
| *+Wh-Subj × +Wh-Obj* | -0.27 | 0.10 | -2.67 | **0.01** |
| *+Wh-Subj × Language<sub>1</sub>* | -0.10 | 0.12 | -0.85 | 0.40 |
| *+Wh-Subj × Language<sub>2</sub>* | 0.26 | 0.10 | 2.73 | **0.01** |
| *+Wh-Obj × Language<sub>1</sub>* | 0.04 | 0.11 | 0.36 | 0.72 |
| *+Wh-Obj × Language<sub>2</sub>* | 0.12 | 0.09 | 1.34 | 0.18 |
| *+Wh-Subj × +Wh-Obj × Lg<sub>1</sub>* | 0.01 | 0.16 | 0.10 | 0.92 |
| *+Wh-Subj × +Wh-Obj × Lg<sub>2</sub>* | **-0.28** | **0.13** | **-2.17** | **0.03** |

Pairwise comparisons revealed that Mandarin Chinese participants were more accurate than English participants (β = 1.02, SE = 0.19, *z*-ratio = 5.44, *p* < 0.001) and Urdu participants (β = 0.72, SE = 0.20, *z*-ratio = 3.61, *p* < 0.01), but there were no differences in accuracy between Urdu and English participants (β = –0.29, SE = 0.19, *z*-ratio = –1.52, *p* = 0.28).

Pairwise comparisons found that English speakers did not respond differently for –Wh-Subject, –Wh-Object trials and –Wh-Subject, +Wh-Object trials (β = –0.04, SE = 0.09, *z*-ratio = –0.4, *p* = 0.70), but were more accurate for +Wh-Subject, –Wh-Object trials compared to +Wh-Subject, +Wh-Object trials (β = 0.52, SE = 0.10, *z*-ratio = 5.03, *p* < 0.001). Similar patterns were observed in Mandarin Chinese. Mandarin Chinese participants did not respond differently for –Wh-Subject, –Wh-Object trials and –Wh-Subject, +Wh-Object trials (β = 0.04, SE = 0.15, *z*-ratio = 0.30, *p* = 0.76), but showed greater accuracy for –Wh-Subject, –Wh-Object trials compared to –Wh-Subject, +Wh-Object trials (β = 0.30, SE = 0.15, *z*-ratio = 2.00, *p* = 0.05). By contrast, Urdu participants responded more accurately for –Wh-Subject, –Wh-Object trials compared to –Wh-Subject, +Wh–Object trials (β = 0.24, SE = 0.12, *z*-ratio = 1.98, *p* = 0.05), and +Wh–Subject, –Wh-Object trials compared to +Wh-Subject, +Wh-Object trials (β = 0.25, SE = 0.13, *z*-ratio = 1.94, *p* = 0.05).

The average presentation time for the critical stimuli across all participants was 206ms (SE = 0.1ms). By language, average presentation times were 208ms (SE < 0.1ms) for English, 203ms (SE < 0.01) for Mandarin Chinese, and 207ms (SE < 0.01ms) for Urdu. Overwhelmingly, due to high accuracy on the task, participants performed at the minimum presentation time, 200ms. However, *t*-tests showed that Mandarin Chinese trial times were significantly faster than English (*t* = –17.43, *p* < 0.01) and Urdu (*t* = –11.73, *p* < 0.01) trial times, and that English trial times were slower than Urdu trial times (*t* = –3.38, *p* < 0.01). Because trial time was dependent on accuracy, the trial time patterns mirror the accuracy results – all 3 language groups performed near ceiling, but Mandarin Chinese-speaking participants outperformed Urdu- and English-speaking participants, and Urdu-speaking participants outperformed English-speaking participants. The significant majority of trials were at the minimum trial time of 200ms (English = 86.5%; Mandarin Chinese = 94.5%; Urdu = 89.4%). The proportion of trial times are presented in Figure 1D.

### 3.2 EEG Analysis

All EEG processing was conducted in MNE-Python (Gramfort et al. 2013), and statistical analysis was conducted in Eelbrain (Brodbeck et al. 2023). Raw EEG data were recorded using FPz as reference, following manufacturer instructions. Raw EEG data were then band-pass filtered off-line using an IIR Butterworth filter from 0.1–40Hz. Flat and noisy channels were then identified, removed, and interpolated. We then re-referenced EEG data to an average reference. Then, independent components analysis (ICA) was used to identify and remove endogenous sources of rhythmic electromagnetic noise, including eye-blinks, ocular movements, and heartbeats. Afterwards, epochs were extracted for the target sentence, from 100ms pre-stimulus onset to 800ms post-stimulus onset, corresponding to the target sentence and the subsequent blank gray screen. Incorrect trials were excluded at this stage. Epochs were baseline corrected to the –100-0ms pre-onset period. Epochs with extreme voltage fluctuations were automatically rejected with a 100μV peak-to-peak threshold, and other bad epochs were rejected after visual inspection. After rejection of incorrect trials and bad epochs, there were 80.4% English trials, 96.2% Mandarin Chinese trials, and 96.8% Urdu trials remaining. For statistical analyses, condition numbers were normalized by subject, and then condition averages were computed.

After pre-processing, the sensor data was analyzed using spatio-temporal cluster-based permutation tests (Maris & Oostenveld 2007). This method allows us to leverage the fact that adjacent time points and sensors are not independent to identify significant clusters in space-time, without the problem of implicit multiple comparisons used in pre-selecting sensors and time windows of analysis. For each time point and sensor, we fit an independent ANOVA to the raw EEG activity across individuals with ±Wh-Subject and ±Wh-Object as factors (μV ∼ ±Wh-Subject × ±Wh-Object). –Wh-Subject and –Wh-Object were coded as 0 and +Wh-Subject and +Wh-Object coded as 1. This produces an *F*-value test-statistic and a *p*-value for each time point and source. Afterwards, points are clustered together if they are adjacent in time and space. This clustering procedure is constrained to clusters of minimum length 20ms, minimum 3 sensors, and must be significant at *p <* 0.05 (English, Mandarin) and *p* < 0.20 (Urdu). All EEG sensors were included in the spatio-temporal clustering procedure, and time windows from 200–800ms were selected. Afterwards, the test-statistics (*F*-values) were summed to produce the cluster-level statistic (‘cluster size’). The ANOVA and clustering procedure was then conducted another 10,000 times, each time randomly permuting the condition labels on the data. These clusters are then ordered by size. This produces a null distribution against which to compare the clusters identified in the observed data. The observed data’s cluster’s *p*-values after correction are their ranks in the bootstrapped null distribution; clusters in the top 5% of this distribution are considered significant at α < 0.05.

A different *p-*value threshold was selected for Urdu due to the smaller sample size. The clustering *p*-value affects cluster size rather than statistical sensitivity. This is because the same *p*-value threshold is used both for the clustering procedure in the observed data and the bootstrapped null distribution.

### 3.4 EEG Results

The EEG results by language are summarized in Figure 1E. The temporal and spatial distribution for each cluster are shown in Figure 2.

**Figure 2.**
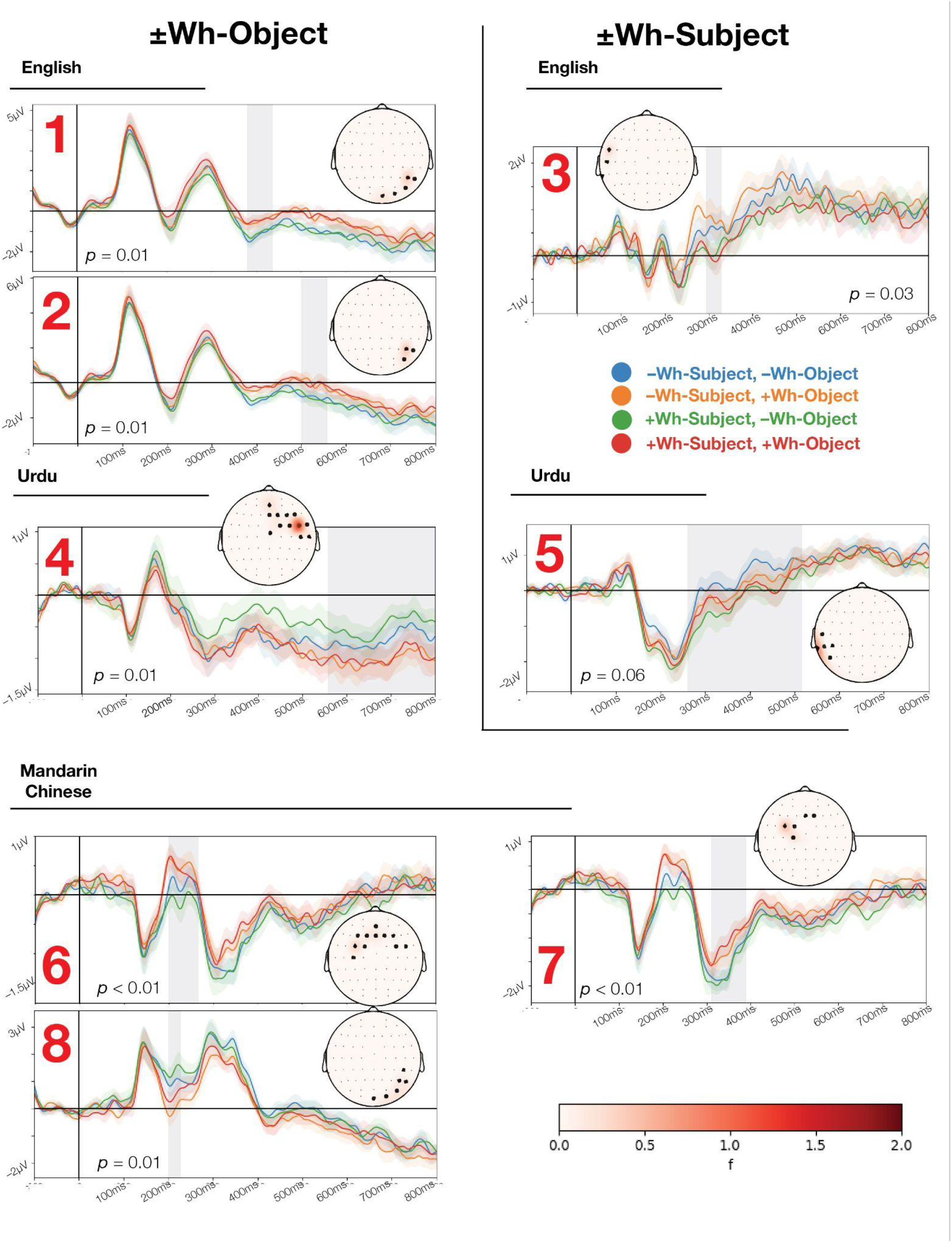
Clusters identified in statistical analysis in English, Urdu, and Mandarin Chinese. Each plot shows the temporal extent of the cluster as an ERP time series, and the spatial extent of the cluster as sensors marked on a topographic plot. Error ribbons in the time series correspond to one standard error above and below the mean at each time point, and gray shading shows the temporal extent of the cluster. Bold black dots in the topographic plot show sensors included in the cluster, and red shading show the *F*-statistic at that point. Reported *p*-values correspond to the position of the observed cluster in the bootstrapped null distribution from the permutation test. Each red number corresponds to the cluster label provided in Section 3.4.

#### English

In English, two ±Wh-Object clusters were observed. The largest cluster was observed from 380–436ms, distributed over right posterior sensors (*p* = 0.01; Figure 2 – Cluster 1). This cluster showed a greater negative peak for –Wh-Objects compared to +Wh-Objects. The second cluster was observed at a later time span, 500–557ms, also over right posterior sensors (*p* = 0.01; Figure 2 – Cluster 2). This cluster showed the same pattern of activity, with a greater sustained negativity for –Wh-Objects compared to +Wh-Objects. A ±Wh-Subject cluster was observed from 294–320ms over left lateral sensors (*p* = 0.03; Figure 2 – Cluster 3). This cluster showed greater positive peak for –Wh-Subjects over +Wh-Subjects. No significant clusters were observed for the ±Wh-Subject × ±Wh-Object interaction.

#### Urdu

In Urdu, one significant cluster was identified for the factor ±Wh-Object. This cluster was observed 558–800ms, centered on central midline and right parietal sensors (*p* = 0.01; Figure 2 – Cluster 4). This cluster showed greater negativity +Wh-Objects compared to –Wh-Objects, although unlike other ±Wh-Object clusters, it was not centered around a clear peak in the waveform. A marginally significant cluster for the factor ±Wh-Subject was observed in left lateral sensors from 261–515ms (*p* = 0.06; Figure 2 – Cluster 5). Like the ±Wh-Object cluster, this cluster showed a sustained negativity for the +Wh-Subject conditions compared to the –Wh-Subject conditions.

#### Mandarin Chinese

In Mandarin Chinese, 3 clusters were identified for factor ±Wh-Object The largest cluster was observed 200–266ms, centered over central midline sensors (*p* < 0.001; Figure 2 – Cluster 6). This cluster showed a greater positivity for +Wh-Objects compared to –Wh-Objects. A second cluster was observed centered over central midline sensors from 311–389ms showing a greater negative peak for +Wh-Objects compared to –Wh-Objects (*p* = 0.01; Figure 2 – Cluster 7). Finally, a third cluster was observed centered over right posterior sensors from 200–227ms, showing greater negativity for +Wh-Objects compared to –Wh-Objects (*p* = 0.01; Figure 2 – Cluster 8). No significant clusters were observed for factor ±Wh-Subject or for the ±Wh-Subject × ±Wh-Object interaction term.

## 4. Discussion

Our research question was whether it is possible to detect a neural response to *wh*-constructions in an RPVP reading task in English, Urdu, and Mandarin Chinese, and whether this response is systematic across the 3 languages. English uses filler-gap dependencies/’movement’ for main clause *wh*-object constructions. These dependencies can trigger complex working memory processes in serial, word-by-word reading or listening. Urdu and Mandarin Chinese, by contrast, use *wh-in-situ* structures, which trigger similarly complex but different working memory retrieval processes, when read word-by-word.

In the behavioral component of the RPVP task, accuracy was overall very high, although lower for English-reading participants. Nonetheless, we observed a systematic reduction in accuracy for *wh*-objects in English, Urdu, and Mandarin Chinese. This was not systematically observed for *wh*-subjects. In all 3 languages, this was also an observed reduction in accuracy for sentences with multiple *wh*-questions specifically, sentences in which both the subject and object are *wh*-phrases are harder than sentences in which only the subject is a *wh*-phrase.

In the EEG recordings, we observed that neural responses in English showed sensitivity to *wh*-subjects around ∼300ms in left lateral sensors, followed by sensitivity to *wh*-vs. non-*wh* objects around ∼400–500ms in right posterior sensors. In Urdu, we found a similar, marginally significant response to *wh*-vs. non-*wh* subjects in left lateral sensors, slightly later than in English, 300–500ms. This was followed by a delayed response to *wh*-objects in midline anterior and right parietal sensors ∼500–800ms. Finally, in Mandarin Chinese, we observed no significant response to *wh*-subjects. However, we observed much more rapid sensitivity to *wh*-objects, with a response ∼200ms in midline anterior and posterior sensors, followed by a distinct response in midline anterior sensors ∼300ms.

Given the variety of differences between English, Urdu, and Mandarin Chinese, it is perhaps not surprising that there was not a single, consistent neural response observed in all 3 languages in both time and space, especially given the nature of the RPVP task. However, some common patterns can be detected. First, neural responses to *wh*-subjects – when detected in English and Urdu – temporally precede neural responses to *wh*-objects. Secondly, this sensitivity to *wh*-subjects appears in left lateral sensors in both English and Urdu. Third, sensitivity to *wh*-objects appears either in midline anterior sensors (Urdu, Mandarin Chinese), or right posterior sensors (English, Mandarin Chinese). Additionally, the first detectable sensitivity to *wh*-elements occurs approximately ∼300ms in English (*wh*-subjects) and Mandarin Chinese (*wh*-objects), with a second ‘late’ effect around ∼500–800ms (English, Urdu). The larger temporal extent of the Urdu cluster also includes this time point, although it is not centered on a clear ‘peak’ like the other cluster time series. These earlier ERP responses to *wh*-elements may be consistent with Krogh & Pylkkänen’s (2024) earliest verb movement findings, also around ∼300ms. Interestingly, none of these neural responses appear to be a typical P600 ERP. The *wh*-object vs. non *wh*-object sensitivity observed 500–800ms in Urdu may have the same approximate timing as a P600 response, but not the usual centro-parietal and posterior distribution on the scalp. Finally, and perhaps most curiously, despite the clear ‘penalty’ for multiple *wh*-constructions in the behavioral tasks, no corresponding interactions were observed in the EEG data, suggesting that the effects of ±Wh-Subject and ±Wh-Object are independent in time and space during processing.

### 4.1 *Wh-in-Situ* and *Wh-*Movement

One surprising finding is that *wh*-objects appear to be more difficult to process than non *wh*-objects, as reflected in the lower accuracy rates in the RPVP behavioral task. This is surprising to observe even in the *wh-in-situ* languages like Urdu and Mandarin Chinese, in which the distinction between the *wh*-objects and non *wh*-objects is not reflected in any other syntactic features of the sentence apart from the selection of a determiner (*wh*-determiners: کونسا/کونسی *kaun sā* / *kaun sī*, 哪 *nǎ*; non *wh*-determiners:,..,_ *yeh*; 这 *zhè*). In the case of Urdu, this may be explained as the increased length of the *wh*-determiner, plus the presence of a determiner-noun agreement in gender required for the *wh*-determiner. However, this could not transparently explain the pattern observed in Mandarin Chinese, in which both determiners are a single character and there is no determiner-noun agreement.. We instead suggest that the explanation for the increased processing cost of the +Wh-Object conditions in all 3 languages reflects the more complex semantic interpretation, i.e., identifying a set of alternatives to the semantic object of the sentence.

In our experiment, we might have expected that the –Wh-Subject, +Wh-Object condition in English may be an ‘outlier’, demonstrating a divergent evoked response. This is the only condition that involves an overt filler-gap dependency in our study. Even in an RPVP task, we might expect this to be particularly difficult to process, since the *wh*-element appears at the left edge of the foveal field, while the verb (or gap) appears at the right edge. Additionally, even with the mitigated demand on memory provided by parallel presentation, a moved *wh*-phrase in English is potentially ambiguous with regard to its thematic role until a relation is established with the verb. Thus, we may have expected an interaction effect in English, in which –Wh-Subject, +Wh-Object (*which book did she read?*) might show a distinct pattern in either time or space compared to +Wh-Subject, +Wh-Object (*who read which book?*). However, we failed to observe this. We only find main effects of ±Wh-Object in English, with no clear distinction between *wh*-objects heading filler-gap dependencies and those that remain *in-situ*.

Nonetheless, we do observe that midline anterior sensors are only included in the ±Wh-Object clusters in Urdu and Mandarin Chinese, i.e., the *wh-in-situ* configurations. It is tempting to conclude that this may reflect some process or representation that is specific to *wh-in-situ* structures. However, if so, it is unclear why we should not observe the same scalp distribution for the ±Wh-Subject clusters, or for the *in-situ wh*-objects in English. We leave the question about clarifying the relationship between the topographic and temporal distribution of our reported effects, the processing mechanics of these structures, and their representational features for future research, as well as identifying their potential neural generators.

### 4.2 *Wh*-Objects vs. *Wh*-Subjects

One surprising finding is that *wh*-subjects do not appear to impact the behavioral responses substantially. This lack of a clear effect of *wh*-subjects may be due to their linear position in the RPVP stimulus. All 3 languages position the subject at the beginning of the clause, thus they are likely to fall outside of the foveal center, and may therefore not be processed as accurately or quickly, which is observed in other RPVP tasks (Snell & Grainger 2019; Dunagan et al. 2024; Flower & Pylkkänen 2024). Moreover, the forms of the *wh*- and non-*wh* subject in all languages and conditions are orthographically short (*wh*-subjects: *who*, کس *kis-ne*, 谁*shéi*; non *wh*-subjects: *she* / *he*; اسنے *us-ne*; 她 / 他 *tā*). This confluence of factors may result in readers not systematically noticing or processing the form of the subject, instead focusing on the verb and the object, which are more central in the field of vision.

However, it is clear that subjects are not completely ignored, as a reliable cluster is observed in the EEG data for English, and marginally in Urdu. Moreover, the presence of *wh*-subject contributes to the multiple *wh* ‘penalty’ observed in the behavioral responses of all 3 languages. Thus, participants cannot completely ignore the syntactic and semantic properties of *wh*-subjects in our task.

One limitation of our design is that we systematically used pronouns and indefinite *wh*-expressions for the subjects, and we used proximate demonstratives and definite or ‘D-linked’ *wh*-determiners for objects (Pesetsky 1987, 2000). Thus, the subject noun phrases never included a lexical noun, whereas the object noun phrases always did. This was a deliberate choice, because we did not want any possible effect of ±Wh-Object or ±Wh-Subject to be explained by the presence or absence of a meaningful, contentful word (i.e., *who* vs. *the* girl; or *what* vs. *the* book). Moreover, pilot testing demonstrated that having two full lexical NPs in both the subject and object position made the RPVP task too difficult for participants to do naturally. Thus, we needed to reduce one of the noun phrases to a simpler form to make the task manageable.

For the object noun phrases, we used both proximate determiners and D-linked *wh*-determiners. In normal usage, these structures presume an established discourse referent for the object. This was done to control for sentence length both between and within languages. For instance, non-D-linked objects in Urdu and Mandarin Chinese typically surface as bare nouns (i.e., کتاب (*kitāb, 书 shū* ‘a book’ or ‘the book’), and thus the *wh*-determiners add a significant number of characters in both languages. This major length confound could render any interpretation of ±Wh-Object effects problematic, especially in an RPVP task. However, D-linked *wh*-constructions are reported to have different processing dynamics than their indefinite counterparts, which may reflect different statuses in memory (Goodall 2004, 2015). The distinctions we report between ±Wh-Subject and ±Wh-Object effects may instead reflect either the different impacts that these noun phrases have on the semantic interpretation of the sentence, their relative prominence or position in the visual signal, their relative statuses in working memory, or a combination of these factors.

One way to address this would be to run a similar set of studies, swapping which noun phrase is the indefinite pronoun and which is a definite / D-linked noun phrase. For instance, in English, we could replicate this study using stimuli like *did this girl read it?, what did this girl read*?, *which girl read it*?, and *which girl read what?*. If we observe a similar set of behavioral and neural responses in an experiment with this design, in which ±Wh-Object effects are later, and more robust in the neural signal, then we have stronger evidence that the grammatical function and syntactic position contribute to the results we report here. By contrast, if we observe that the pattern of results ‘flip’, such that ±Wh-Subject effects follow ±Wh-Object effects, and have a more reliable effect on the neural signal, then this would support the hypothesis that the form and semantics of these noun phrases determined the effects observed in these studies.

Lastly, what explains the apparent ‘penalty’ to multiple *wh*-constructions, and why would this be observed only in the behavioral data? Multiple *wh*-constructions are reported to require a ‘pair-list’ reading (Dayal 1996), in which an answer consists of a list of pair-wise answers to each *wh*-question, (19). The +Wh-Subject, +Wh-Object conditions in our study are the only items which require this more complex semantic representation, and thus may be more difficult to interpret. Moreover, detecting that this interpretation is necessarily involves computing a semantic structure that involves computing the alternatives over both the subject and the object. Perceptually, this also requires that participants notice that both of the verb’s arguments are *wh*-phrases. If so, then this observed penalty in the behavioral response may correspond to integration of the *wh*-subject and *wh*-object, and computation of the more complex semantics. If this is the right interpretation, this demonstrates sensitivity to more complex semantic structure and relations between grammatical elements. This demonstrates that readers in an RPVP task are computing graammatical features that are not associated with any particular morpheme or word.

### (19)

a. Question: ‘Who read this book?’ Answers: { *John* read this book, *Jane* read this book…}
b. Question: ‘Who read which book?’ Answers: { [ ‘*John* read *Carrie, Jane* read *Christine*…’], [‘*Jane* read *Carrie, John* read *Christine*…’], …}

### 4.3 RPVP and Long-Distance Relations

Finally, what do these results say about the processes deployed in parallel reading? Previous authors have stressed the mind’s capacity to extract the semantic ‘gist’ of a sentence from a quick glance. Asano et al. (2011) showed that readers can recover a Japanese sentence displayed for a short span more accurately if the sentence is pragmatically likely and coherent, and Massol et a. (2021) reported similar findings in French. To explain these kinds of facts, Wen et al. (2019) argue for an interactive process in which grammatical cues interact with lexical access to facilitate the extraction of some possible meanings or analyses of the sentence. This is evidenced by the finding that ungrammatical, scrambled sentences are much harder in an RPVP task, and elicit distinct neural responses early in EEG and MEG recordings, often around ∼200–300ms (Snell & Grainger 2019; Fallon & Pylkkänen 2024; Dunagan et al. 2024; Dufau et al. 2024), compatible with the view that some grammatical information can be detected with rapid presentation.

However, it is less clear what level of detail this grammatical ‘sketch’ may have. In an MEG experiment, Flower & Pylkkänen (2024) found that earlier brain responses are not sensitive to transposition of adjacent words in sentences presented in an RPVP paradigm, but later brain responses are. Dunagan et al. (2024) found a lack of sensitivity to subject-verb agreement violations in both behavioral and neural responses in English (see also Fallon & Pylkkänen 2024). However, they demonstrate distinct neural responses ∼150ms to sentences with singular and plural subjects. This suggests that readers can identify the morphosyntactic properties of the subject, i.e. they are not ‘blind’ to the presence or absence of inflectional morphology. But, they do not integrate this into a larger model of the structure of the sentence. A similar result is shown by Khokhar et al. (2024), in which case marking and verb gender-marking elicit distinct responses in Urdu, but Urdu readers do not exhibit sensitivity to ungrammatical subject-verb agreement or object-verb agreement. Taken together, it is tempting to infer that some grammatical relations, such as argument-verb agreement, may not be fully ‘noticed’ or fully processed in an RPVP paradigm. Instead, readers might only seek to identify one or two key nouns and verbs that are relevant to guessing the likely meaning of the sentence. This may be enough to extract the ‘gist,’ the who-did-what-to-whom, without computing an entire syntactic and semantic structure. Put differently, the finding that grammatical vs. scrambled sentences elicit different early neural responses may correspond to how easy it is to identify a likely subject nouns, a likely verb, or a likely object noun to reconstruct the most plausible meaning of the sentence. In English, the agent is usually a noun somewhere on the left, and the object is usually a noun somewhere on the right, etc. These findings are arguably also compatible with the early insensitivity to local transpositions reported by Flower & Pylkkänen (2024).

However, our results show that this is too simplistic. Readers do, in fact, extract some grammatical and relational information, beyond the identity of a few contentful lexical items. Indeed, the clear distinction between *wh*-objects and non *wh*-objects in Urdu and Mandarin show that early neural responses and off-line behavioral responses are affected by a change in a single determiner, a part of the ‘functional’/grammatical vocabulary of the language. Moreover, as argued in Section 4.2, the sensitivity to multiple *wh*-constructions in all 3 languages may indicate computation of semantic relations between elements in the sentence. Thus, we argue that readers of English, Urdu, and Mandarin Chinese can extract key elements of grammatical dependencies above and beyond basic nouns and verbs. The difference between the findings reported here for *wh*-constructions and those reported for argument-verb constructions reported in English (Dunagan et al. 2024; Fallon & Pylkkänen 2024) and Urdu (Khokhar et al. 2024) may reflect the fact that the relations between *wh*-constructions are vital for identifying the logical semantics of the sentence, whereas argument-verb agreement may be less relevant.

## 5. Conclusion

How is grammatical structure represented in the brain? This is a challenging question to answer, in part because similar structures across languages may place very different demands on memory, depending on superficial features such as word order or writing system, which require different psycholinguistic computations to be executed at different times. Nonetheless, the organization of the language network in the brain appears to be largely invariant across languages, and linguistic theory has long argued that grammatical phenomena rely on shared representational features across languages. How can we relate the universal to the language-particular?

Here, we leverage the RPVP paradigm, in which short sentences of different grammatical structures are displayed for a short duration. We suggest that this paradigm may allow us to sidestep the differences in processing dynamics that would normally be engendered by serial word-by-word reading or listening. Instead of measuring the brain’s evoked response to a word given the prior context and parsing state, the RPVP paradigm allows us to measure the brain’s evoked response to the *sentence*. We propose that this provides a new way for integrating insights from comparative syntax, the systematic study of language similarities and differences, to the cognitive neuroscience of language. We propose that we can make advances in understanding how the brain represents particular classes of grammatical structures by systematic comparison between languages using paradigms like RPVP that minimize the differences in real-time processing.

The mechanisms supporting reading entire sentences rapidly presented are still under debate (e.g., Wen et al. 2019; Dufau et al. 2024; Fallon & Pylkkänen 2024; Flower & Pylkkänen 2024; Dunagan et al. 2024). Nonetheless, we showed that readers can extract key grammatical information from sentences displayed in 200ms in English, Urdu, and Mandarin Chinese. Moreover, we show that neural and behavioral responses in an RPVP task are sensitive to *wh*-constructions, even if the superficial reification of this construction is the choice of a determiner (i.e., *wh*-object vs. non *wh*-objects in Mandarin Chinese). Thus, the processes deployed when a reader encounters an entire sentence for 200ms are sensitive to ‘functional’/grammatical vocabulary items like *wh*-pronouns and *wh*-determiners (see also Flower & Pylkkänen 2024), as observed both in the neural responses and the behavioral responses.

Lastly, with specific regard to *wh*-constructions, our results show essentially three basic patterns. Neural responses to *wh*-objects surface either in right posterior or midline anterior sensors, and either from ∼200–400ms or ∼500–800ms. Neural responses to *wh*-subjects surface in left lateral sensors around ∼200-300ms, or not at all. What remains to be clarified is what features of the language determine where and when these responses are observed. Lastly, the off-line behavioral responses across all 3 languages showed that *wh*-objects are harder to process than their non *wh*-alternatives, even if there is no filler-gap dependency involved, and that multiple *wh*-constructions also engender processing difficulty. We hope that future work can elaborate on the processing mechanics deployed during reading in RPVP tasks, grammatical properties, and the spatio-temporal distribution of the evoked neural responses observed in this study.

## Acknowledgments

We thank Nigel Flower and Shulin Zhang for assistance with creating Mandarin Chinese stimuli and Maria Ali for assistance with creating the Urdu stimuli and recruiting Urdu participants. We also thank Liina Pylkkänen for comments and discussion on this project as part of a larger collaboration. Finally, we thank John Hale for feedback and suggestions.

This work was partially supported by UGA Seed funding awarded to Dustin A. Chacón, and NSF Grant 1903783 awarded to John Hale.

